# Identification of expression quantitative trait loci in the skeletal muscle of Thoroughbreds reveals heritable variation in expression of genes relevant to cofactor metabolism

**DOI:** 10.1101/713669

**Authors:** G. Farries, K. Bryan, C. L. McGivney, P. A. McGettigan, K. F. Gough, J.A. Browne, D. E. MacHugh, L. M. Katz, E. W. Hill

## Abstract

**Summary:** While over ten thousand genetic loci have been associated with phenotypic traits and inherited diseases in genome-wide association studies, in most cases only a relatively small proportion of the trait heritability is explained and biological mechanisms underpinning these traits have not been clearly identified. Expression quantitative trait loci (eQTL) are subsets of genomic loci shown experimentally to influence gene expression. Since gene expression is one of the primary determinants of phenotype, the identification of eQTL may reveal biologically relevant loci and provide functional links between genomic variants, gene expression and ultimately phenotype. Skeletal muscle (gluteus medius) gene expression was quantified by RNA-seq for 111 Thoroughbreds (47 male, 64 female) in race training at a single training establishment sampled at two time-points: at rest (*n* = 92) and four hours after high-intensity exercise (*n* = 77); *n* = 60 were sampled at both time points. Genotypes were generated from the Illumina Equine SNP70 BeadChip. Applying a Benjamini-Hochberg corrected *P*-value threshold (*P*_FDR_ < 0.05), association tests identified 3,583 *cis*-eQTL associated with expression of 1,456 genes at rest; 4,992 *cis*-eQTL associated with the expression of 1,922 genes post-exercise; 1,703 *trans*-eQTL associated with 563 genes at rest; and 1,219 *trans*-eQTL associated with 425 genes post-exercise. The gene with the highest *cis*-eQTL association at both time-points was the endosome-associated-trafficking regulator 1 gene (*ENTR1*; Rest: *P*_FDR_ = 3.81 × 10^−27^, Post-exercise: *P*_FDR_ = 1.66 × 10^−24^), which has a potential role in the transcriptional regulation of the solute carrier family 2 member 1 glucose transporter protein (SLC2A1). Functional analysis of genes with significant eQTL revealed significant enrichment for cofactor metabolic processes. These results suggest heritable variation in genomic elements such as regulatory sequences (e.g. gene promoters, enhancers, silencers), microRNA and transcription factor genes, which are associated with metabolic function and may have roles in determining end-point muscle and athletic performance phenotypes in Thoroughbred horses.

## Introduction

In the 6,000 years since horses were first domesticated on the Eurasian steppe, there has been strong artificial selection for various athletic traits (Levine, 1999). Selection for athleticism is perhaps most clearly manifested in the Thoroughbred, which has undergone over 300 years of intense selection for speed and racing performance. As a result the Thoroughbred has a highly developed musculature, with a skeletal muscle mass ∼10% greater than other horse breeds (∼55% compared to ∼42%) (Gunn, 1987), accompanied by decreased body fat (Kearns et al., 2002), superior glycogen storage capacity (Votion et al., 2012), increased mitochondrial volume (compared to other mammals) (Kayar et al., 1989) and a high degree of plasticity in skeletal muscle fibre composition (Rivero, 2004).

The response of equine skeletal muscle to training has been well studied. These responses in general increase the oxidative capacity of the muscle, such as fibre type switching from fast-twitch glycolytic fibres to slow-twitch, high-oxidative fibres (Serrano et al., 2000), an increase in oxidative phosphorylation (Votion et al., 2012) and increased mitochondrial volume (Tyler et al., 1998). Training also elicits an increase in skeletal muscle mass (Rivero et al., 1996), mediated through hyperplastic growth as opposed to marked hypertrophy (Rivero et al., 1996; Rivero et al., 2002).

The transcriptional response to exercise and training in skeletal muscle has been studied in the Thoroughbred (McGivney et al., 2009; Eivers et al., 2010; McGivney et al., 2010; Eivers et al., 2012; Bryan et al., 2017). Initially, reverse transcription quantitative real-time polymerase chain reaction (RT-qPCR) was used to quantify expression of 18 candidate genes in response to a standardised exercise test on a high-speed treadmill (Eivers et al., 2010). Significant differential expression of *CKM, COX4I1, COX4I2, PDK4, PPARG1A* and *SLC2A4* four hours post-exercise was detected. The availability of a dedicated equine microarray allowed gene expression to be measured across 9,333 expressed sequence tags (ESTs). This technology was then used to examine the changes in gene expression induced by exercise, without *a priori* knowledge of the genes involved (McGivney et al., 2009). Analysis of the differentially expressed genes showed a functional enrichment of genes involved in insulin signalling, focal adhesion, hypertrophic and apoptotic pathways. Digital gene expression was used to investigate the transcriptional response to a ten-month training protocol (McGivney et al., 2010), identifying functional enrichment of genes relevant to aerobic metabolism. More recently, RNA sequencing (RNA-seq) was used to investigate the response to both exercise and training, and a network biology approach was employed to identify relevant functional modules that highlighted the role of autophagy (Bryan et al., 2017) While these studies provide insight to the genes involved in the transcriptional response to exercise, they do not reveal whether there is variation in the transcriptional response among individuals and how this may influence skeletal muscle function.

Expression quantitative trait loci (eQTL) are genomic variants, typically SNPs, that are associated with variation in RNA transcript abundance. Jansen and Nap (2001) introduced the concept of ‘genetical genomics’ where genomic loci were associated with cellular intermediates, such as transcript abundance, to catalogue functional relevance for non-coding variants. This was a particularly important development because the clear majority (>85%) of QTLs detected in genome-wide association (GWA) studies are located in non-coding regions (Hindorff et al., 2009; Brown et al., 2013).

*Cis*-eQTL are genetic variants that alter gene expression in an allele-specific manner and are typically located in gene regulatory regions (Wittkopp, 2005; Westra and Franke, 2014). Identification of true *cis*-eQTL requires aligning reads to their chromosome of origin; consequently, many studies have by convention defined any eQTLs within 1 Mb of the transcription start site (TSS) of the gene they act on as *cis*. Conversely, *trans*-eQTL act in a less direct manner, altering the expression of a secondary genome product—for example, a transcription factor or a microRNA—that regulates expression of a distant gene elsewhere in the genome (Wittkopp, 2005).

The study of eQTL in skeletal muscle to-date has been largely to investigate functional variants in the pathogenesis of type II diabetes (T2D) in humans (Mason et al., 2011; Keildson et al., 2014b; Sajuthi et al., 2016b). While GWA studies for T2D have identified loci associated with disease risk, these studies have not provided information on the function of these variants or the mechanism by which they contribute to disease. Keildson et al. (2014b) performed an eQTL investigation using skeletal muscle biopsies from 104 human subjects and identified an association between the rs4547172 SNP and muscle phosphofructokinase gene (*PFKM*) expression. Furthermore, the study found that increased expression of *PFKM* was associated with increased resting plasma insulin (an endophenotype) and T2D (an end-point phenotype). This example shows that an eQTL approach can identify functional links between genomic variants, gene expression, endophenotypes and ultimately, disease.

Variation in gene expression has been found to be highly heritable (Monks et al., 2004; Stranger et al., 2007; Wright et al., 2014). Given the influence of gene expression on phenotype, detection of heritable variation in skeletal muscle gene expression may provide insight into genomic loci contributing to variation in exercise and performance related phenotypes.

In this study, we hypothesised that there is heritable variation in the Thoroughbred skeletal muscle transcriptional response to exercise and training, and that this variation may have implications for athletic performance.

## Methods

### Ethics statement

University College Dublin Animal Research Ethics Committee approval (AREC-P-12-55-Hill), a licence from the Department of Health (B100/3525) and informed owner consent were obtained.

### Cohort

Skeletal muscle biopsy samples (gluteus medius) were collected from 111 horses (47 male, 64 female) born between 2011 and 2012. All horses were based at a single training yard, under the supervision of a single trainer and under similar management and feeding regimes. The 111 horses used for the study were produced from 19 different sires and 94 different dams.

Biopsies were collected at two time points: untrained at rest (UR) and untrained four hours post-exercise (UE). Of the 111 horses, 60 were sampled at both time points. In total 92 UR samples and 77 UE samples were collected. The horses were defined as untrained because they had completed ≤ four sprint exercise bouts (e.g., work days) prior to sampling.

### Exercise test

The exercise stimulus was an intense sprint bout of exercise (work day). Horses were initially walked on an automated horse walker for 30-60 min, followed by 5-10 min of walking in hand. Under saddle there was an initial warm-up period of 300 m walk and 700 m of trot and slow canter down the incline of the track. The work day was performed on a 1,500 m all-weather woodchip gallop track, with the final 800 m straight set on a 2.7% incline. The sprint portion of the exercise bout consisted of the horses galloping at high intensity for 800-1,000 m up the incline of the gallop.

For 34/77 UE horses, whole blood was collected at rest and five minutes post-exercise into fluoride oxalate tubes. Samples were centrifuged, and plasma lactate concentrations measured on-site using a YSI2300 STAT PLUS auto analyser (YSI UK Ltd, Hampshire, UK). These measurements were used to validate the intensity of the exercise test performed.

### Biopsy sampling

Percutaneous needle muscle biopsies (approximately 300 mg) were obtained from the ventral compartment of the middle gluteal muscle using the method described by Valette et al. (1999). All UR samples were collected between 7:30 am and 11:30 am. UE samples were taken four hours after completion of the exercise test, as this has previously been shown to be a timepoint where the greatest change in gene expression in response to acute exercise was observed (McGivney et al., 2009; Eivers et al., 2010). Muscle samples were stored in RNAlater (Thermo Fisher, Massachusetts, USA) for 24 hours at 4°C then stored at −20°C prior to RNA extraction.

### RNA extraction and quality control

Total RNA was extracted from approximately 70 mg tissue using a protocol combining TRIzol reagent (Thermo Fisher), DNase I treatment (Qiagen, Hilden, Germany) and an RNeasy Mini-Kit (Qiagen). RNA was quantified using a Nano Drop ND1000 spectrophotometer V 3.5.27 (Thermo Fisher). RNA quality was assessed using the RNA integrity number (RIN) on an Agilent Bioanalyser with the RNA 6000 Nano LabChip kit6 (Agilent, Cork, Ireland).

### RNA sequencing

Indexed, strand-specific Illumina sequencing libraries were prepared using the TruSeq Stranded mRNA Library Preparation Kit LT (Illumina, San Diego, USA). Libraries were pooled with 18-20 indexed libraries per pool and sequenced on an Illumina HiSeq 2500 using a Rapid Run flow cell and reagents to generate 100 bp paired-end reads. Each pool was sequenced across both lanes of the flow cell (dual lane loading). Demultiplexed sequence data was then converted to FASTQ format. Sequencing was performed by the Research Technology Support Facility, Michigan State University.

### RNA-seq data workflow

Quality control of the sequence reads was performed using *FastQC* [version: 0.11.5] (Andrews, 2010). *STAR* aligner [version: 2.5.2b] (Dobin et al., 2013) was used to map reads to the Equine reference genome EquCab2 (Ensembl release 62). After mapping, *featureCounts* [version: 1.5.0] was used to assign reads to genes (Liao et al., 2014). Data for each sample from each sequencing lane was then merged where concordance was >99% between lanes. Count data was quantile normalised using *preprocessCore* [version: 1.40.0] (Bolstad, 2017) within the R environment [version: 3.5.1] (R Core Team, 2017), and the log_2_ of quantile-normalised count data calculated.

### Genotyping

Genomic DNA was extracted from whole blood using the Maxwell 16 automated DNA purification system (Promega, Madison, WI). Horses were genotyped on the Illumina Equine SNP70 BeadChip (Illumina). A genetic versus phenotypic sex check was performed. SNPs with a genotyping rate of <95%, and individuals with a genotyping rate <95% were excluded. SNPs with a minor allele frequency (MAF) <0.10 were removed. The remaining 43,988 SNPs were then pruned based on pairwise linkage disequilibrium (LD) using a sliding window with an LD threshold of *r*^2^ > 0.7, a window size of 50, and a step of 5 in *PLINK* [version: 1.09] (Chang et al., 2015). A set of 15,995 SNPs were used for the eQTL analysis.

### eQTL analysis

eQTL were determined using a linear model within *matrixEQTL* [version: 2.1.1] (Shabalin, 2012); including sex, age at sampling (days) and RNA-seq batch as covariates. Tests of association were corrected using the Benjamini-Hochberg procedure (Benjamini and Hochberg, 1995) and eQTL with a corrected *P*-value (*P*_FDR_) < 0.05 were catalogued for UR and UE samples separately. eQTL located within 1 Mb of the transcription start site (TSS) of the gene they were associated were designated as *cis*, and those located >1Mb from the TSS were designated *trans*. Significant results were then compared against genes previously identified in the skeletal muscle transcriptional response to acute, high-intensity exercise (a work day; 3,241 genes) and transcriptional response to a six-month period of training (3,405 genes) (Bryan et al., 2017).

### Functional enrichment analysis

Genes with significant eQTL were investigated for enrichment of biological processes using gene ontology (GO) categories (Ashburner et al., 2000) with the *clusterProfiler* package [version: 3.10.1] (Yu et al., 2012) within the R environment. Equine Ensembl IDs were mapped to annotated human orthologs, retrieved from the BioMart database (Kasprzyk, 2011) and GO enrichment performed using the annotation from the human genome annotation package *org.Hs.eg.db* [version: 2.12.0] (Carlson, 2017). The background gene set was the complement of genes expressed in skeletal muscle identified in this study (13,384 genes; 12,707 mapped to human orthologs). A threshold for significant enrichment was set at <0.05 after adjustment using the Benjamini-Hochberg procedure (*P*_FDR_) (Benjamini and Hochberg, 1995). The number of genes assigned to each Biological Process (Gene count) and proportion of genes associated with that cluster out of all the genes expressed (Gene ratio) were also reported. Results were visualised using the *clusterProfiler* package (Yu et al., 2012).

## Results

### Cohort

UR horses had a mean age of 611.7 days (range: 513-787 days), UE horses had a mean age of 757.5 days (range: 617-1,283 days). Dates of commencing preparatory training were available for 90 of the UR horses; 21 of the UR horses were sampled prior to breaking, 69 were sampled after breaking with a mean of 41.5 days after commencing preparatory training (range: 5-154 days) (Table 1). UE horses were sampled on average 156.6 days after commencing preparatory training (range: 31-307). UR horses had an average of 41.5 days submaximal training (range: 5-154) and UE had on average 48.6 (range: 19-152) (Table 1). UR horses had completed a mean of 0.3 work days (range: 0-4), UE horses completed a mean of 0.5 WDs prior to sampling (range: 0-3) (Table 2). A subset of 34 of the UE horses had a mean peak post-exercise plasma lactate concentration of 28.2 mmol/L, and a mean resting plasma concentration of 0.4 mmol/L. All RNA samples used for RNA-seq had a RIN greater than 7.0, the UR cohort had a mean RIN of 8.0 (range: 7.2−9.3) and the UE cohort had a mean RIN of 8.1 (range: 7.0−9.3).

**Table 1.**
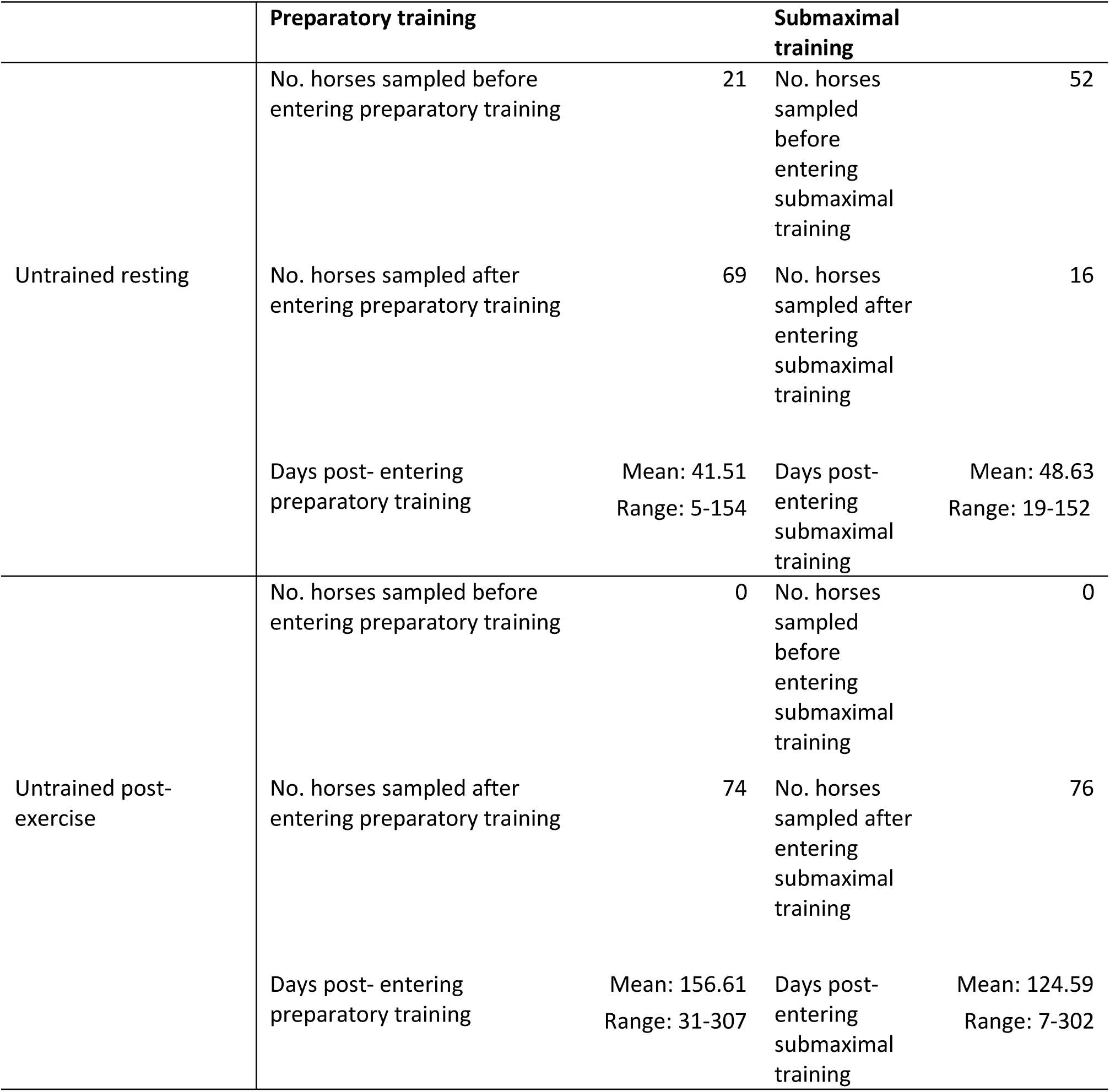
Description of prior training completed by individuals in the untrained resting and untrained post-exercise cohorts.

|  | Preparatory training |  | Submaximal training |  |
| --- | --- | --- | --- | --- |
| Untrained resting | No. horses sampled before entering preparatory training | 21 | No. horses sampled before entering submaximal training | 52 |
|  | No. horses sampled after entering preparatory training | 69 | No. horses sampled after entering submaximal training | 16 |
|  | Days post- entering preparatory training | Mean: 41.51<br>Range: 5-154 | Days post- entering submaximal training | Mean: 48.63<br>Range: 19-152 |
| Untrained post-exercise | No. horses sampled before entering preparatory training | 0 | No. horses sampled before entering submaximal training | 0 |
|  | No. horses sampled after entering preparatory training | 74 | No. horses sampled after entering submaximal training | 76 |
|  | Days post- entering preparatory training | Mean: 156.61<br>Range: 31-307 | Days post- entering submaximal training | Mean: 124.59<br>Range: 7-302 |

**Table 2.**
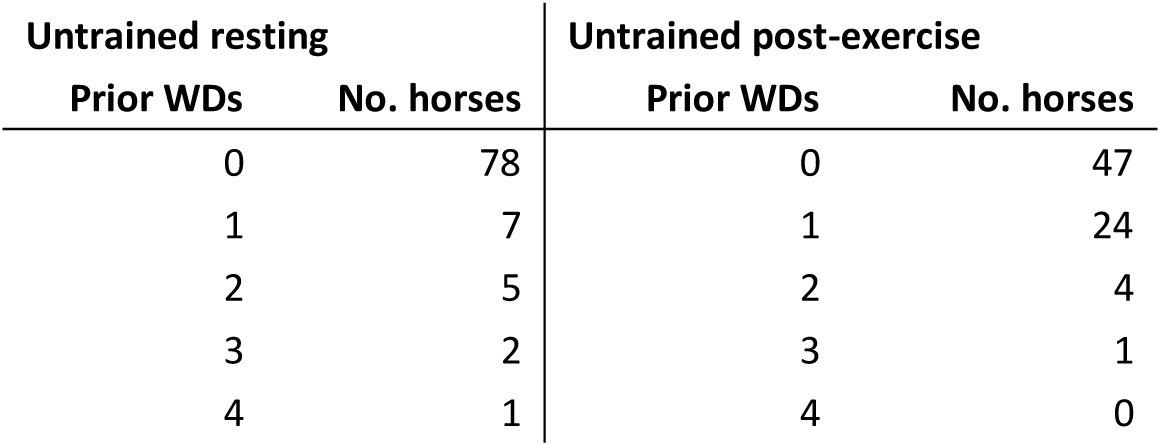
Number of prior high-intensity sprint bouts (work days, WDs) completed prior to sampling for horses within untrained resting and untrained post-exercise cohorts.

| Untrained resting |  | Untrained post-exercise |  |
| --- | --- | --- | --- |
| Prior WDs | No. horses | Prior WDs | No. horses |
| 0 | 78 | 0 | 47 |
| 1 | 7 | 1 | 24 |
| 2 | 5 | 2 | 4 |
| 3 | 2 | 3 | 1 |
| 4 | 1 | 4 | 0 |

### eQTL discovery

Using the full complement of 13,384 genes, 3,582 *cis*-eQTL and 1,703 *trans*-eQTL were detected in UR samples (*P*_FDR_ < 0.05). The 3,582 *cis*-eQTL were associated with expression of 1,456 genes. The gene with the strongest *cis*-eQTL (BIEC2-707785) in UR horses was the endosome associated trafficking regulator 1 gene (*ENTR1*; *P*_FDR_ = 3.81 × 10^−27^) (Figure S2, Table 3). GO enrichment analysis of the *cis* regulated genes in UR samples showed that the most significantly enriched Biological Process was ‘GO:0006805 xenobiotic metabolicprocess ‘ (*P*_FDR_ = 3.02 × 10^−7^, Gene Ratio = 33/1,614). ‘GO:0051186 cofactor metabolic process’ (*P*_*FDR*_ = 1.42 × 10^−4^) was also significantly enriched and had the largest Gene Ratio (105/1,614) (Figure 1, Table S1).

**Table 3.**
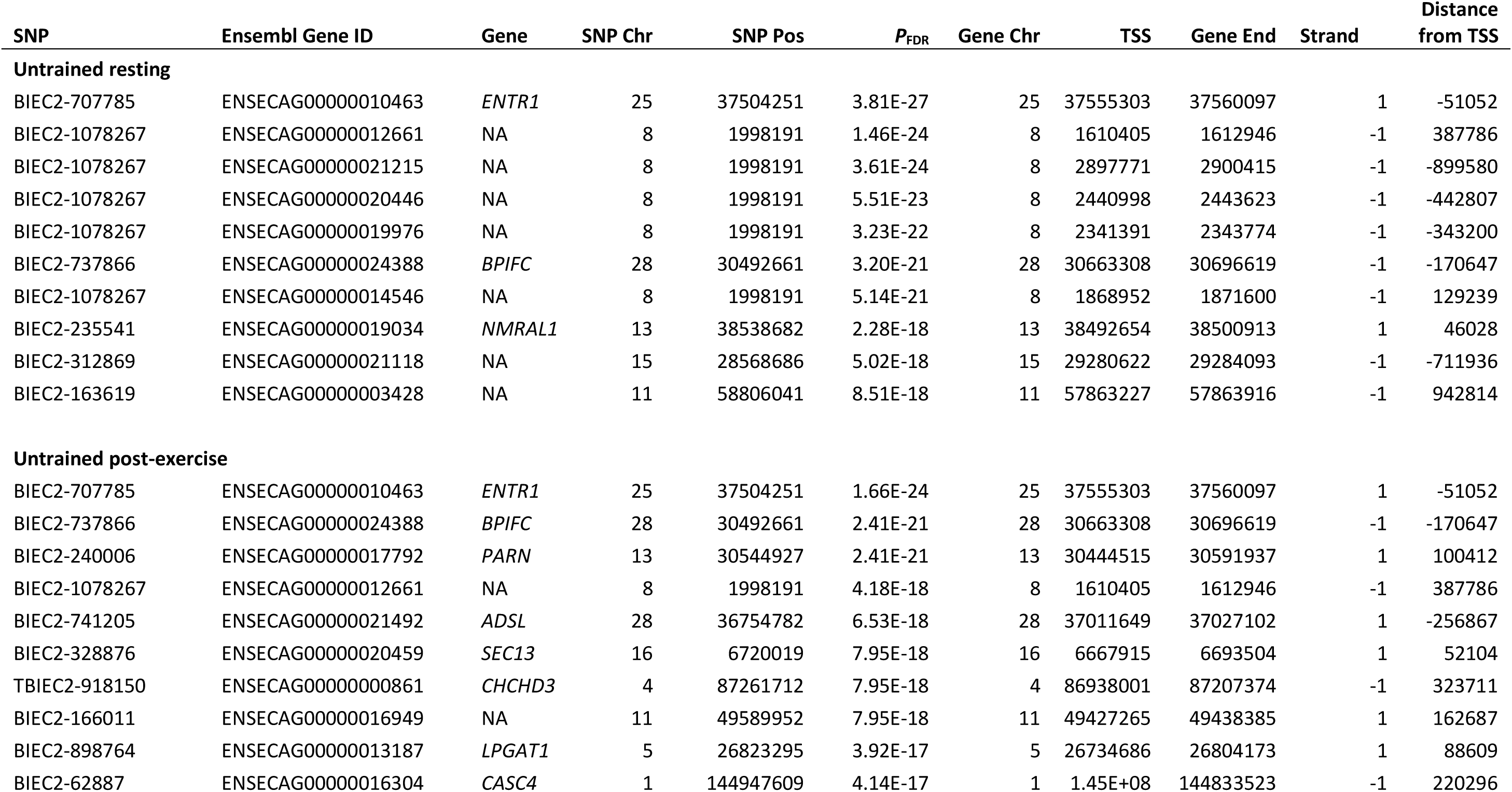
top ten genes by strongest *cis*-eQTL association.

| SNP | Ensembl Gene ID | Gene | SNP Chr | SNP Pos | *P*<sub>FDR</sub> | Gene Chr | TSS | Gene End | Strand | Distance from TSS |
| --- | --- | --- | --- | --- | --- | --- | --- | --- | --- | --- |
| **Untrained resting** |  |  |  |  |  |  |  |  |  |  |
| BIEC2-707785 | ENSECAG00000010463 | *ENTR1* | 25 | 37504251 | 3.81E-27 | 25 | 37555303 | 37560097 | 1 | -51052 |
| BIEC2-1078267 | ENSECAG00000012661 | NA | 8 | 1998191 | 1.46E-24 | 8 | 1610405 | 1612946 | -1 | 387786 |
| BIEC2-1078267 | ENSECAG00000021215 | NA | 8 | 1998191 | 3.61E-24 | 8 | 2897771 | 2900415 | -1 | -899580 |
| BIEC2-1078267 | ENSECAG00000020446 | NA | 8 | 1998191 | 5.51E-23 | 8 | 2440998 | 2443623 | -1 | -442807 |
| BIEC2-1078267 | ENSECAG00000019976 | NA | 8 | 1998191 | 3.23E-22 | 8 | 2341391 | 2343774 | -1 | -343200 |
| BIEC2-737866 | ENSECAG00000024388 | *BPIFC* | 28 | 30492661 | 3.20E-21 | 28 | 30663308 | 30696619 | -1 | -170647 |
| BIEC2-1078267 | ENSECAG00000014546 | NA | 8 | 1998191 | 5.14E-21 | 8 | 1868952 | 1871600 | -1 | 129239 |
| BIEC2-235541 | ENSECAG00000019034 | *NMRAL1* | 13 | 38538682 | 2.28E-18 | 13 | 38492654 | 38500913 | 1 | 46028 |
| BIEC2-312869 | ENSECAG00000021118 | NA | 15 | 28568686 | 5.02E-18 | 15 | 29280622 | 29284093 | -1 | -711936 |
| BIEC2-163619 | ENSECAG00000003428 | NA | 11 | 58806041 | 8.51E-18 | 11 | 57863227 | 57863916 | -1 | 942814 |
| **Untrained post-exercise** |  |  |  |  |  |  |  |  |  |  |
| BIEC2-707785 | ENSECAG00000010463 | *ENTR1* | 25 | 37504251 | 1.66E-24 | 25 | 37555303 | 37560097 | 1 | -51052 |
| BIEC2-737866 | ENSECAG00000024388 | *BPIFC* | 28 | 30492661 | 2.41E-21 | 28 | 30663308 | 30696619 | -1 | -170647 |
| BIEC2-240006 | ENSECAG00000017792 | *PARN* | 13 | 30544927 | 2.41E-21 | 13 | 30444515 | 30591937 | 1 | 100412 |
| BIEC2-1078267 | ENSECAG00000012661 | NA | 8 | 1998191 | 4.18E-18 | 8 | 1610405 | 1612946 | -1 | 387786 |
| BIEC2-741205 | ENSECAG00000021492 | *ADSL* | 28 | 36754782 | 6.53E-18 | 28 | 37011649 | 37027102 | 1 | -256867 |
| BIEC2-328876 | ENSECAG00000020459 | *SEC13* | 16 | 6720019 | 7.95E-18 | 16 | 6667915 | 6693504 | 1 | 52104 |
| TBIEC2-918150 | ENSECAG00000000861 | *CHCHD3* | 4 | 87261712 | 7.95E-18 | 4 | 86938001 | 87207374 | -1 | 323711 |
| BIEC2-166011 | ENSECAG00000016949 | NA | 11 | 49589952 | 7.95E-18 | 11 | 49427265 | 49438385 | 1 | 162687 |
| BIEC2-898764 | ENSECAG00000013187 | *LPGAT1* | 5 | 26823295 | 3.92E-17 | 5 | 26734686 | 26804173 | 1 | 88609 |
| BIEC2-62887 | ENSECAG00000016304 | *CASC4* | 1 | 144947609 | 4.14E-17 | 1 | 1.45E+08 | 144833523 | -1 | 220296 |

**Figure 1.**
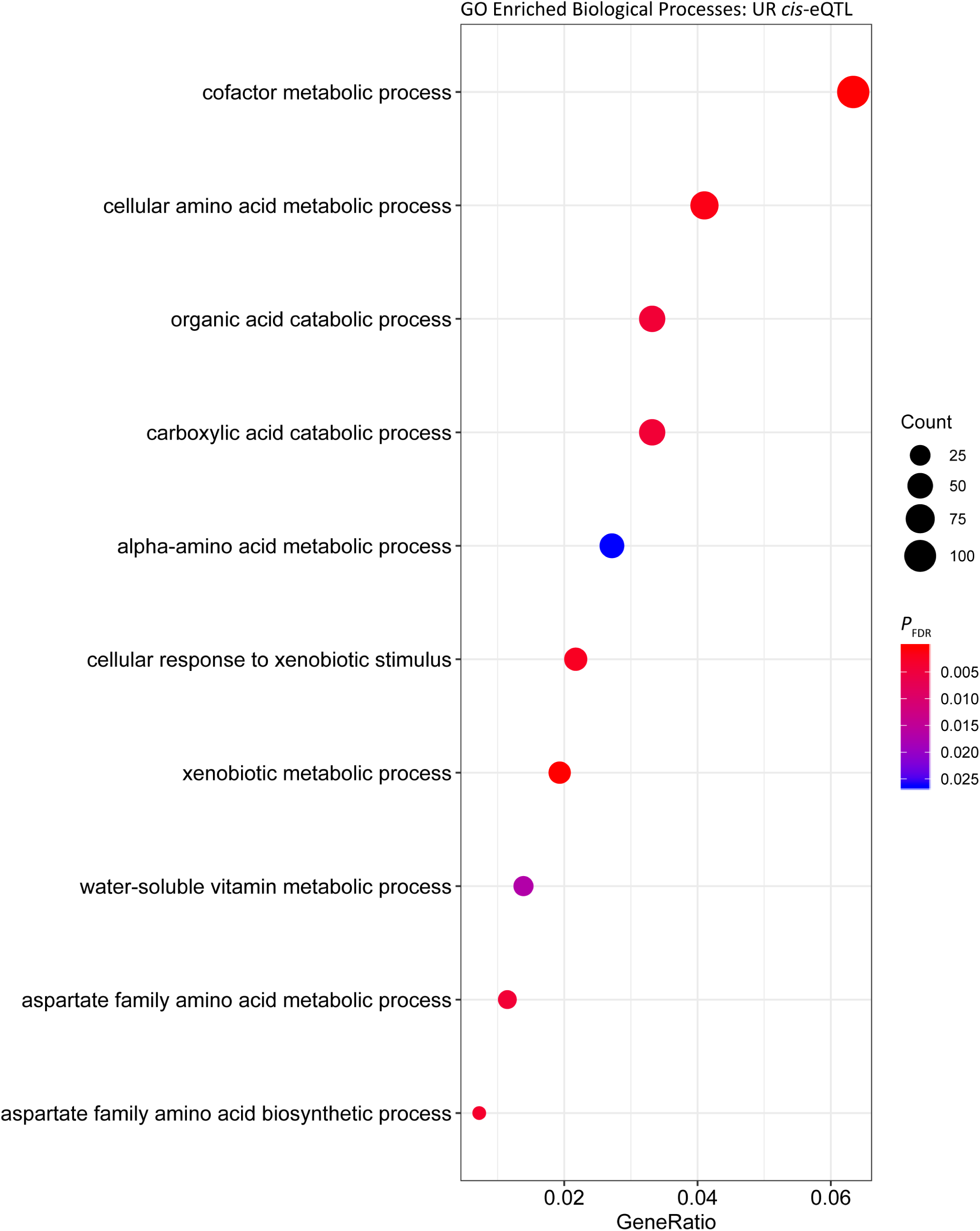
Gene ontology enrichment of biological processes for genes under *cis* regulation in the untrained resting cohort.

In the UR cohort 1,219 *trans*-eQTL were associated with 425 genes. The majority 70.39% (858) were located on the same chromosome as the associated gene, and 29.61% (361) were associated with genes located on different chromosomes. The most significant *trans*-eQTL was BIEC2-526896 on ECA20 and expression of the DEAH-box helicase 16 gene (*DHX16*) also located on ECA20 1.49 Mb downstream from BIEC2-526896 (*P*_FDR_ = 3.50 × 10^−17^)(Table 4). Functional analysis of the *trans* eGenes showed enrichment of ‘interferon-gamma-mediated signalling’ (*P*_FDR_ = 6.06 × 10^−4^, Gene Ratio = 13/340) (Figure 2, Table S2). The functional categories with the highest Gene Ratios were ‘cofactor metabolic process’ (*P*_*FDR*_ = 0.01, Gene Ratio = 29/340) and ‘monocarboxylic acid metabolic process’ (*P*_*FDR*_ = 0.02, Gene Ratio = 29/340) (Figure 2, Table S2).

**Table 4.** top ten genes by strongest *trans*-eQTL association.

| SNP | Ensembl Gene ID | Gene | SNP Chr | SNP Pos | $P_{FDR}$ | Gene Chr | TSS | Gene End | Strand | Distance from TSS |
| --- | --- | --- | --- | --- | --- | --- | --- | --- | --- | --- |
| **Untrained resting** |  |  |  |  |  |  |  |  |  |  |
| BIEC2-526896 | ENSECAG00000008537 | *DHX16* | 20 | 28206968 | 3.50E-17 | 20 | 29693915 | 29708818 | -1 | -1486947 |
| BIEC2-218753 | ENSECAG00000015585 | *TBL2* | 13 | 9657509 | 8.69E-14 | 13 | 11241845 | 11246395 | -1 | -1584336 |
| BIEC2-526896 | ENSECAG00000015782 | NA | 20 | 28206968 | 5.20E-13 | 20 | 29364710 | 29368260 | 1 | -1157742 |
| BIEC2-526896 | ENSECAG00000015505 | NA | 20 | 28206968 | 3.57E-12 | 20 | 30173637 | 30178981 | -1 | -1966669 |
| BIEC2-554291 | ENSECAG00000019318 | NA | 20 | 29116874 | 3.57E-12 | 20 | 32450503 | 32452484 | -1 | -3333629 |
| BIEC2-991035 | ENSECAG00000012956 | *DCPS* | 7 | 31604226 | 9.10E-12 | 7 | 35117723 | 35153555 | 1 | -3513497 |
| BIEC2-526896 | ENSECAG00000021750 | NA | 20 | 28206968 | 2.27E-11 | 20 | 31193941 | 31197235 | -1 | -2986973 |
| BIEC2-526791 | ENSECAG00000009368 | NA | 20 | 27782384 | 2.27E-11 | 20 | 29049395 | 29052907 | 1 | -1267011 |
| BIEC2-554291 | ENSECAG00000021750 | NA | 20 | 29116874 | 4.38E-11 | 20 | 31193941 | 31197235 | -1 | -2077067 |
| BIEC2-31048 | ENSECAG00000024546 | NA | 1 | 73056328 | 5.71E-11 | 1 | 70638960 | 70656636 | -1 | 2417368 |
| **Untrained post-exercise** |  |  |  |  |  |  |  |  |  |  |
| UKUL2765 | ENSECAG00000010213 | *MCCC2* | 14 | 93893549 | 1.12E-13 | 14 | 92304084 | 92460487 | -1 | 1589465 |
| BIEC2-3702 | ENSECAG00000004989 | NA | 1 | 9825107 | 4.38E-12 | 1 | 96257111 | 96258450 | 1 | -86432004 |
| BIEC2-991035 | ENSECAG00000012956 | *DCPS* | 7 | 31604226 | 2.03E-11 | 7 | 35117723 | 35153555 | 1 | -3513497 |
| BIEC2-526896 | ENSECAG00000015505 | NA | 20 | 28206968 | 6.37E-11 | 20 | 30173637 | 30178981 | -1 | -1966669 |
| BIEC2-210079 | ENSECAG00000012497 | *LIMK1* | 13 | 10604233 | 6.76E-11 | 13 | 11610902 | 11634978 | 1 | -1006669 |
| BIEC2-554291 | ENSECAG00000019318 | NA | 20 | 29116874 | 2.91E-10 | 20 | 32450503 | 32452484 | -1 | -3333629 |
| BIEC2-526896 | ENSECAG00000015782 | NA | 20 | 28206968 | 4.06E-10 | 20 | 29364710 | 29368260 | 1 | -1157742 |
| BIEC2-554291 | ENSECAG00000015505 | NA | 20 | 29116874 | 8.06E-10 | 20 | 30173637 | 30178981 | -1 | -1056763 |
| BIEC2-63245 | ENSECAG00000016304 | *CASC4* | 1 | 1.48E+08 | 1.04E-09 | 1 | 1.45E+08 | 1.45E+08 | -1 | 3045045 |
| BIEC2-3702 | ENSECAG00000009972 | *LHPP* | 1 | 9825107 | 2.16E-09 | 1 | 8372229 | 8511110 | -1 | 1452878 |

**Figure 2.**
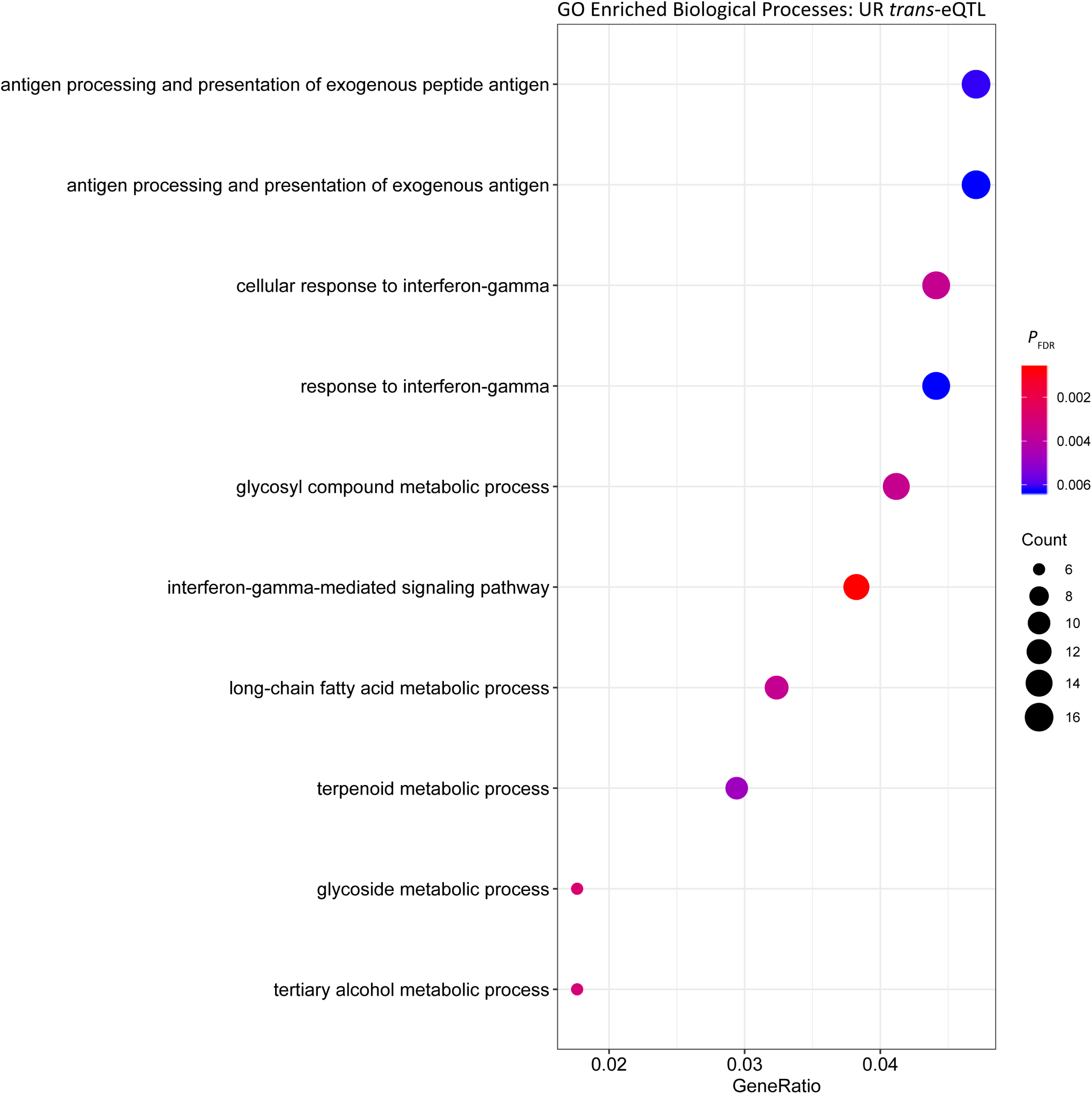
Gene ontology enrichment of biological processes for genes under *trans* regulation in the untrained resting cohort.

In the UE cohort 4,992 *cis*-eQTL were associated with the expression of 1,922 genes. The most significant *cis*-eQTL was BIEC2-707785 on ECA25 and *ENTR1* (*P*_FDR_ = 1.66 × 10^−24^), as was the case in the UR cohort (Table 3). The strongest *trans*-eQTL association was between BIEC2-165011 on ECA11 and transcript ENSECAG00000016949 (*P*_FDR_ = 1.12 × 10^−13^) (Table 4). Similar to UR samples, the majority (75.45%; 544) of *trans*-eQTL were located on the same chromosome and 24.55% (177) were on different chromosomes.

Analysis of the *cis* regulated genes in UE samples showed that similar to the UR cohort, the most significantly enriched Biological Process was ‘cofactor metabolic process’ (*P*_FDR_ = 6.40 × 10^−7^, Gene Ratio = 112/1,579) (Figure 3, Table S3). Comparable results were obtained for enrichment of Biological Processes among putative *trans* regulated genes, with ‘cofactor metabolic process’ the most significantly enriched (*P*_FDR_ = 5.06 × 10^−7^, Gene Ratio = 33/235).

**Figure 3.**
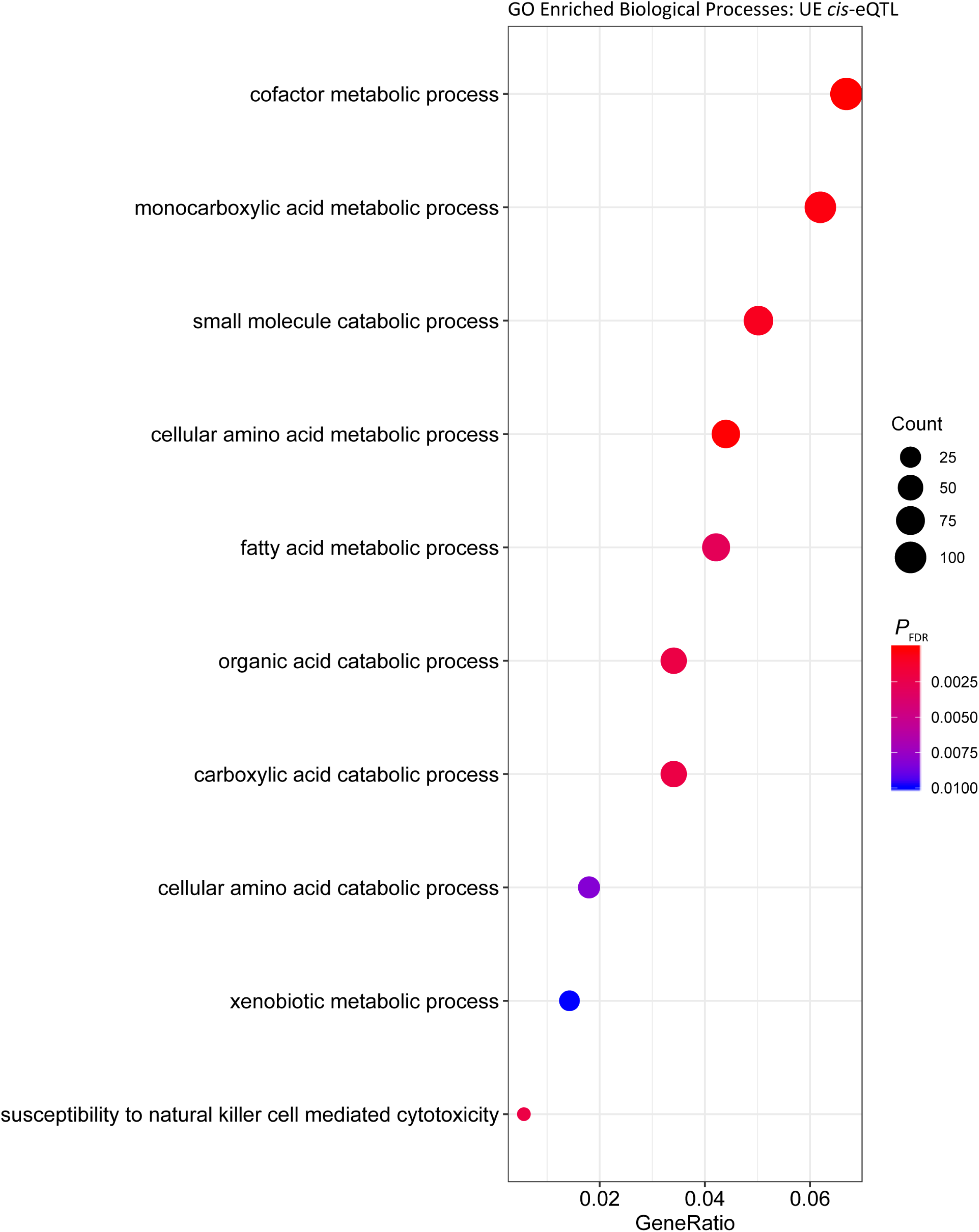
Gene ontology enrichment of biological processes for genes under *cis* regulation in the post-exercise cohort.

**Figure 4.**
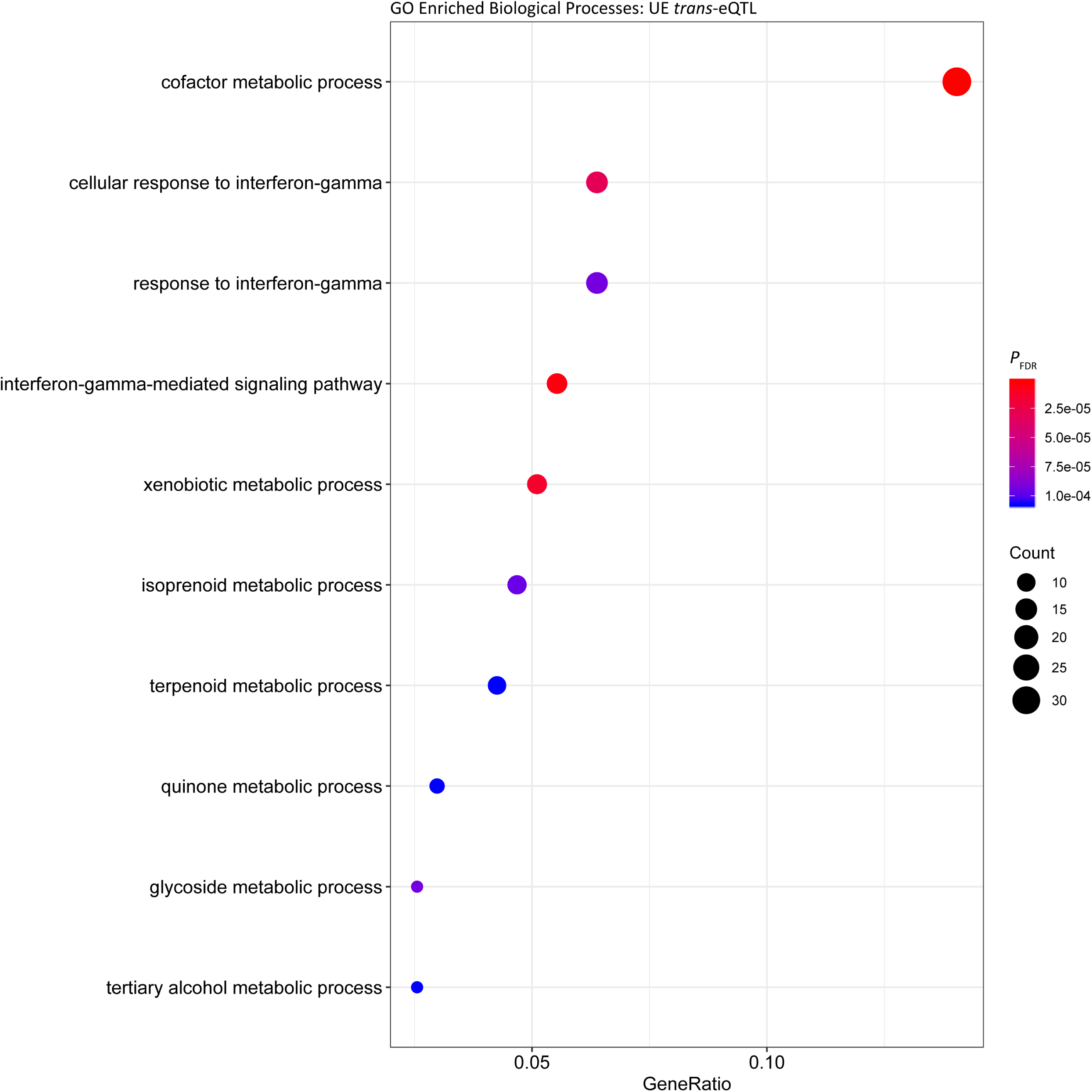
Gene ontology enrichment of biological processes for genes under *trans* regulation in the post-exercise cohort.

### Genetic regulation of exercise relevant genes

The total set of genes with expression changes associated with eQTLs (i.e. eGenes) were queried against genes that we have reported from the same dataset to be differentially expressed post-exercise (Bryan et al., 2017). Of the 3,582 UR *cis*-eQTL, 913 were associated with genes differentially expressed in response to exercise. The most significant association was between BIEC2-285235 and the CCR4-NOT transcription complex subunit 11 gene (*CNOT11*; *P*_FDR_ = 3.00 × 10^−15^) (Table 5). Of the 1,703 UR *trans*-eQTL, 144 were associated with exercise relevant genes. The most significant *trans*-eQTL was between BIEC2-1061469 and the TAL bHLH transcription factor 2 gene (*TAL2*; *P*_FDR_ = 3.03 × 10^−10^) (Table 6).

**Table 5.** top ten *cis*-eQTL identified in genes differentially expressed in response to exercise.

| SNP | Ensembl Gene ID | Gene | SNP Chr | SNP Pos | $P_{FDR}$ | Gene Chr | TSS | Gene End | Strand | Distance from TSS |
| --- | --- | --- | --- | --- | --- | --- | --- | --- | --- | --- |
| **Untrained resting** |  |  |  |  |  |  |  |  |  |  |
| BIEC2-285235 | ENSECAG00000011967 | *CNOT11* | 15 | 8238549 | 3.00E-15 | 15 | 8266315 | 8276804 | -1 | -27766 |
| BIEC2-240006 | ENSECAG00000017792 | *PARN* | 13 | 30544927 | 4.42E-15 | 13 | 30444515 | 30591937 | 1 | 100412 |
| BIEC2-209001 | ENSECAG00000019190 | *IFT22* | 13 | 9212034 | 5.33E-15 | 13 | 9228359 | 9233728 | -1 | -16325 |
| BIEC2-1035081 | ENSECAG00000008723 | NA | 7 | 14212942 | 5.82E-14 | 7 | 14574017 | 14585790 | -1 | -361075 |
| BIEC2-830969 | ENSECAG00000012898 | *PNLDC1* | 31 | 1778298 | 3.95E-13 | 31 | 2140239 | 2157442 | 1 | -361941 |
| BIEC2-872079 | ENSECAG00000024368 | *IPO9* | 30 | 28911273 | 1.23E-12 | 30 | 28818157 | 28849624 | 1 | 93116 |
| BIEC2-988215 | ENSECAG00000018541 | *C2CD2L* | 7 | 26800468 | 4.48E-12 | 7 | 26815981 | 26823699 | 1 | -15513 |
| BIEC2-911228 | ENSECAG00000024971 | *GSTM4* | 5 | 57817054 | 4.38E-11 | 5 | 58455048 | 58459646 | -1 | -637994 |
| BIEC2-609981 | ENSECAG00000011021 | *NDUFAF5* | 22 | 9904659 | 7.88E-11 | 22 | 10479889 | 10512792 | -1 | -575230 |
| TBIEC2-57788 | ENSECAG00000013327 | *SLC24A1* | 1 | 1.27E+08 | 9.47E-11 | 1 | 126764822 | 1.27E+08 | -1 | -3157 |
| **Untrained post-exercise** |  |  |  |  |  |  |  |  |  |  |
| BIEC2-240006 | ENSECAG00000017792 | *PARN* | 13 | 30544927 | 2.41E-21 | 13 | 30444515 | 30591937 | 1 | 100412 |
| BIEC2-209001 | ENSECAG00000019190 | *IFT22* | 13 | 9212034 | 5.64E-13 | 13 | 9228359 | 9233728 | -1 | -16325 |
| BIEC2-1082617 | ENSECAG00000008479 | *SDR16C5* | 9 | 27686807 | 3.39E-12 | 9 | 27293107 | 27307849 | 1 | 393700 |
| BIEC2-1035081 | ENSECAG00000008723 | NA | 7 | 14212942 | 6.06E-12 | 7 | 14574017 | 14585790 | -1 | -361075 |
| TBIEC2-513760 | ENSECAG00000000973 | NA | 2 | 73037320 | 1.15E-11 | 2 | 72462318 | 72935365 | 1 | 575002 |
| BIEC2-770455 | ENSECAG00000011697 | *TMCC3* | 28 | 19829349 | 2.82E-11 | 28 | 19803131 | 19811873 | -1 | 26218 |
| BIEC2-14401 | ENSECAG00000010786 | *PGAM1* | 1 | 32009184 | 6.25E-11 | 1 | 31823570 | 31825848 | -1 | 185614 |
| BIEC2-412279 | ENSECAG00000023566 | *DHRS9* | 18 | 48009567 | 1.04E-10 | 18 | 48741376 | 48752360 | 1 | -731809 |
| BIEC2-1029754 | ENSECAG00000019879 | *ISCU* | 8 | 11582493 | 2.78E-10 | 8 | 12361828 | 12395086 | -1 | -779335 |
| BIEC2-909217 | ENSECAG00000009353 | *AGMO* | 4 | 48124305 | 3.15E-10 | 4 | 47940706 | 48126316 | -1 | 183599 |

**Table 6.** top ten *trans*-eQTL identified in genes differentially expressed in response to exercise.

| SNP | Ensembl Gene ID | Gene | SNP Chr | SNP Pos | $P_{FDR}$ | Gene Chr | TSS | Gene End | Strand | Distance from TSS |
| --- | --- | --- | --- | --- | --- | --- | --- | --- | --- | --- |
| **Untrained resting** |  |  |  |  |  |  |  |  |  |  |
| BIEC2-658237 | ENSECAG00000004995 | TAL2 | 25 | 9914109 | 3.03E-10 | 25 | 11725340 | 11725720 | 1 | -1811231 |
| BIEC2-53419 | ENSECAG00000013327 | SLC24A1 | 1 | 124846027 | 3.72E-08 | 1 | 126764822 | 126790389 | -1 | -1918795 |
| BIEC2-58303 | ENSECAG00000013327 | SLC24A1 | 1 | 128180460 | 2.14E-06 | 1 | 126764822 | 126790389 | -1 | 1415638 |
| BIEC2-913863 | ENSECAG00000024971 | GSTM4 | 5 | 63531114 | 5.08E-06 | 5 | 58455048 | 58459646 | -1 | 5076066 |
| BIEC2-959879 | ENSECAG00000024971 | GSTM4 | 5 | 61035410 | 3.24E-05 | 5 | 58455048 | 58459646 | -1 | 2580362 |
| BIEC2-496328 | ENSECAG00000014441 | ZNF330 | 2 | 89041634 | 3.72E-05 | 2 | 90360628 | 90375365 | -1 | -1318994 |
| BIEC2-311700 | ENSECAG00000017760 | ELK3 | 15 | 53973327 | 5.26E-05 | 28 | 21110553 | 21148433 | 1 | NA |
| BIEC2-907346 | ENSECAG00000018012 | SELENBP1 | 5 | 44309385 | 6.82E-05 | 5 | 45898879 | 45906904 | 1 | -1589494 |
| BIEC2-59128 | ENSECAG00000013327 | SLC24A1 | 1 | 129427658 | 1.09E-04 | 1 | 126764822 | 126790389 | -1 | 2662836 |
| BIEC2-60026 | ENSECAG00000013327 | SLC24A1 | 1 | 130739790 | 1.59E-04 | 1 | 126764822 | 126790389 | -1 | 3974968 |
| **Untrained post-exercise** |  |  |  |  |  |  |  |  |  |  |
| BIEC2-1053404 | ENSECAG00000011541 | PRDX2 | 7 | 51698577 | 1.51E-08 | 7 | 45573319 | 45575352 | 1 | 6125258 |
| BIEC2-996678 | ENSECAG00000011541 | PRDX2 | 7 | 41340513 | 2.58E-08 | 7 | 45573319 | 45575352 | 1 | -4232806 |
| BIEC2-658237 | ENSECAG00000004995 | TAL2 | 25 | 9914109 | 9.79E-08 | 25 | 11725340 | 11725720 | 1 | -1811231 |
| BIEC2-1028898 | ENSECAG00000019879 | ISCU | 8 | 10602506 | 1.29E-06 | 8 | 12361828 | 12395086 | -1 | -1759322 |
| BIEC2-1028319 | ENSECAG00000019879 | ISCU | 8 | 10149470 | 9.98E-06 | 8 | 12361828 | 12395086 | -1 | -2212358 |
| BIEC2-1051774 | ENSECAG00000023096 | TPGS2 | 8 | 51698367 | 2.02E-05 | 8 | 56940922 | 56994571 | -1 | -5242555 |
| BIEC2-420441 | ENSECAG00000009547 | WDR12 | 18 | 76611180 | 4.93E-05 | 18 | 77655352 | 77674750 | -1 | -1044172 |
| TBIEC2-1144261 | ENSECAG00000008479 | SDR16C5 | 9 | 30312051 | 4.94E-05 | 9 | 27293107 | 27307849 | 1 | 3018944 |
| TBIEC2-1149509 | ENSECAG00000008479 | SDR16C5 | 9 | 40285361 | 6.34E-05 | 9 | 27293107 | 27307849 | 1 | 12992254 |
| BIEC2-1052388 | ENSECAG00000023096 | TPGS2 | 8 | 53636613 | 1.07E-04 | 8 | 56940922 | 56994571 | -1 | -3304309 |

Within the UE cohort 4,992 *cis*-eQTL were identified, 1,132 of which were associated with genes differentially expressed post-exercise. The strongest association was between BIEC2-240006 and the polyA-specific ribonuclease gene (*PARN*; *P*_FDR_ = 2.41 × 10^−21^) (Table 5). Of the UR trans-eQTL, 121 eQTL were associated with eGenes in the transcriptional exercise response. The strongest *trans*-eQTL in the UE cohort was BIEC2-1053404, associated with expression of the peroxiredoxin 2 gene (*PRDX2*; *P*_FDR_ = 1.51 × 10^−8^) (Table 6).

### Genetic regulation of training relevant genes

Using 3,405 genes that were differentially expressed in response to training (Bryan et al, 2017), we examined our results based on eQTL associated with genes within this transcriptional response. Within the UR cohort, 609 of the 3,582 *cis*-eQTL were associated with training response genes. The strongest association was between BIEC2-1061469 and the spindle and expression of the kinetochore associated complex subunit 1 gene (*SKA1*; *P*_FDR_ = 9.80 × 10^−18^) (Table 7). Of the 1,703 UR *trans*-eQTL, 145 were associated with training response genes. The most significant association was between BIEC2-658237 and *TAL2* (*P*_FDR_ = 3.03 × 10^−10^) (Table 8).

**Table 7.** top *cis*-eQTL identified in genes differentially expressed in response to training.

| SNP | Ensembl Gene ID | Gene | SNP Chr | SNP Pos | $P_{FDR}$ | Gene Chr | TSS | Gene End | Strand | Distance from TSS |
| --- | --- | --- | --- | --- | --- | --- | --- | --- | --- | --- |
| **Untrained resting** |  |  |  |  |  |  |  |  |  |  |
| BIEC2-1061469 | ENSECAG00000023484 | SKA1 | 8 | 69117228 | 9.80E-18 | 8 | 68814638 | 68828326 | 1 | 302590 |
| BIEC2-1061072 | ENSECAG00000023484 | SKA1 | 8 | 68498709 | 3.57E-17 | 8 | 68814638 | 68828326 | 1 | -315929 |
| TBIEC2-1028240 | ENSECAG00000022151 | MATK | 7 | 2020098 | 5.03E-13 | 7 | 2272120 | 2277087 | -1 | -252022 |
| BIEC2-974177 | ENSECAG00000022151 | MATK | 7 | 1546395 | 1.12E-12 | 7 | 2272120 | 2277087 | -1 | -725725 |
| BIEC2-872079 | ENSECAG00000024368 | IPO9 | 30 | 28911273 | 1.23E-12 | 30 | 28818157 | 28849624 | 1 | 93116 |
| BIEC2-988215 | ENSECAG00000018541 | C2CD2L | 7 | 26800468 | 4.48E-12 | 7 | 26815981 | 26823699 | 1 | -15513 |
| BIEC2-472999 | ENSECAG00000021368 | FGGY | 2 | 732213 | 7.94E-12 | 2 | 549904 | 834638 | -1 | 182309 |
| BIEC2-472598 | ENSECAG00000015014 | ACADL | 2 | 248866 | 1.08E-11 | 2 | 124580 | 165039 | -1 | 124286 |
| BIEC2-609981 | ENSECAG00000011021 | NDUFAF5 | 22 | 9904659 | 7.88E-11 | 22 | 10479889 | 10512792 | -1 | -575230 |
| TBIEC2-57788 | ENSECAG00000013327 | SLC24A1 | 1 | 1.27E+08 | 9.47E-11 | 1 | 1.27E+08 | 1.27E+08 | -1 | -3157 |
| **Untrained post-exercise** |  |  |  |  |  |  |  |  |  |  |
| UKUL3712 | ENSECAG00000010475 | IL33 | 23 | 27442828 | 8.07E-16 | 23 | 27442768 | 27454175 | 1 | 60 |
| BIEC2-974177 | ENSECAG00000022151 | MATK | 7 | 1546395 | 1.96E-14 | 7 | 2272120 | 2277087 | -1 | -725725 |
| TBIEC2-1028240 | ENSECAG00000022151 | MATK | 7 | 2020098 | 7.95E-14 | 7 | 2272120 | 2277087 | -1 | -252022 |
| BIEC2-1082617 | ENSECAG00000008479 | SDR16C5 | 9 | 27686807 | 3.39E-12 | 9 | 27293107 | 27307849 | 1 | 393700 |
| BIEC2-472598 | ENSECAG00000015014 | ACADL | 2 | 248866 | 8.29E-12 | 2 | 124580 | 165039 | -1 | 124286 |
| TBIEC2-513760 | ENSECAG00000000973 | NA | 2 | 73037320 | 1.15E-11 | 2 | 72462318 | 72935365 | 1 | 575002 |
| BIEC2-360074 | ENSECAG00000014129 | TMEM42 | 16 | 42029863 | 2.23E-10 | 16 | 41642464 | 41643652 | -1 | 387399 |
| BIEC2-472999 | ENSECAG00000021368 | FGGY | 2 | 732213 | 2.74E-10 | 2 | 549904 | 834638 | -1 | 182309 |
| BIEC2-1029754 | ENSECAG00000019879 | ISCU | 8 | 11582493 | 2.78E-10 | 8 | 12361828 | 12395086 | -1 | -779335 |
| BIEC2-136659 | ENSECAG00000023268 | CALHM5 | 10 | 64788579 | 3.77E-10 | 10 | 65017017 | 65021618 | 1 | -228438 |

**Table 8.** top *trans*-eQTL identified in genes differentially expressed in response to training.

| SNP | Ensembl Gene ID | Gene | SNP Chr | SNP Pos | $P_{FDR}$ | Gene Chr | TSS | Gene End | Strand | Distance from TSS |
| --- | --- | --- | --- | --- | --- | --- | --- | --- | --- | --- |
| **Untrained resting** |  |  |  |  |  |  |  |  |  |  |
| BIEC2-658237 | ENSECAG00000004995 | TAL2 | 25 | 9914109 | 3.03E-10 | 25 | 11725340 | 11725720 | 1 | -1811231 |
| BIEC2-1118794 | ENSECAG00000023484 | SKA1 | 8 | 67640813 | 4.93E-10 | 8 | 68814638 | 68828326 | 1 | -1173825 |
| BIEC2-53419 | ENSECAG00000013327 | SLC24A1 | 1 | 124846027 | 3.72E-08 | 1 | 126764822 | 126790389 | -1 | -1918795 |
| BIEC2-58303 | ENSECAG00000013327 | SLC24A1 | 1 | 128180460 | 2.14E-06 | 1 | 126764822 | 126790389 | -1 | 1415638 |
| BIEC2-59128 | ENSECAG00000013327 | SLC24A1 | 1 | 129427658 | 1.09E-04 | 1 | 126764822 | 126790389 | -1 | 2662836 |
| BIEC2-196080 | ENSECAG00000017776 | PITPNM1 | 12 | 25661692 | 1.28E-04 | 12 | 27228025 | 27239903 | -1 | -1566333 |
| BIEC2-542299 | ENSECAG00000021368 | FGGY | 20 | 7321704 | 1.34E-04 | 2 | 549904 | 834638 | -1 | NA |
| BIEC2-60026 | ENSECAG00000013327 | SLC24A1 | 1 | 130739790 | 1.59E-04 | 1 | 126764822 | 126790389 | -1 | 3974968 |
| BIEC2-580450 | ENSECAG00000011021 | NDUFAF5 | 22 | 9444819 | 2.10E-04 | 22 | 10479889 | 10512792 | -1 | -1035070 |
| BIEC2-328903 | ENSECAG00000020701 | CAV3 | 16 | 6817902 | 2.43E-04 | 16 | 8001463 | 8011813 | -1 | -1183561 |
| **Untrained post-exercise** |  |  |  |  |  |  |  |  |  |  |
| BIEC2-1053404 | ENSECAG00000011541 | PRDX2 | 7 | 51698577 | 1.51E-08 | 7 | 45573319 | 45575352 | 1 | 6125258 |
| BIEC2-996678 | ENSECAG00000011541 | PRDX2 | 7 | 41340513 | 2.58E-08 | 7 | 45573319 | 45575352 | 1 | -4232806 |
| BIEC2-658237 | ENSECAG00000004995 | TAL2 | 25 | 9914109 | 9.79E-08 | 25 | 11725340 | 11725720 | 1 | -1811231 |
| BIEC2-1028898 | ENSECAG00000019879 | ISCU | 8 | 10602506 | 1.29E-06 | 8 | 12361828 | 12395086 | -1 | -1759322 |
| BIEC2-344350 | ENSECAG00000014129 | TMEM42 | 16 | 43032493 | 4.09E-06 | 16 | 41642464 | 41643652 | -1 | 1390029 |
| BIEC2-1028319 | ENSECAG00000019879 | ISCU | 8 | 10149470 | 9.98E-06 | 8 | 12361828 | 12395086 | -1 | -2212358 |
| BIEC2-130408 | ENSECAG00000023268 | CALHM5 | 10 | 67512852 | 2.28E-05 | 10 | 65017017 | 65021618 | 1 | 2495835 |
| TBIEC2-1144261 | ENSECAG00000008479 | SDR16C5 | 9 | 30312051 | 4.94E-05 | 9 | 27293107 | 27307849 | 1 | 3018944 |
| BIEC2-809036 | ENSECAG00000005431 | CLRN2 | 3 | 107506494 | 6.34E-05 | 3 | 106028106 | 106037626 | -1 | 1478388 |
| TBIEC2-1149509 | ENSECAG00000008479 | SDR16C5 | 9 | 40285361 | 6.34E-05 | 9 | 27293107 | 27307849 | 1 | 12992254 |

Within the UE cohort 766 of the 4,992 *cis*-eQTL were associated with training response genes. The most significant *cis*-eQTL association was between UKUL3712 and the interleukin 33 gene (*IL33*; *P*_FDR_ = 8.07 × 10^−16^) (Table 7). Of the 1,219 UE *trans*-eQTL 90 were associated with genes relevant to training. As with the exercise relevant genes, the strongest UE *trans*-eQTL was between BIEC2-1053404 and *PRDX2* (*P*_FDR_ = 1.51 × 10^−8^) (Table 8).

## Discussion

Using a systems genetics approach we have integrated RNA-seq and genome-wide SNP data for a large cohort of Thoroughbred horses in active race training that were maintained in a single environment. This strategy has allowed us to detect significant *cis* and *trans* eQTL in equine skeletal muscle that are likely to be relevant to an exercise phenotype, adaptation to training, that is one of the hallmarks of the Thoroughbred. A total of 4,992 *cis*-eQTL associated with the expression of 1,922 distinct genes were identified in the UR cohort; and 4,886 *cis*-eQTL associated with the expression of 1,875 genes were identified in the UE cohort. Fewer *trans*-eQTL were detected (UR: 1,703; UE: 1,219), which is consistent with previous studies, and likely due to the greater statistical power required to identify *trans*-eQTL (Westra and Franke, 2014).

The gene with the most significant association with a *cis*-eQTL in the UR and UE cohorts was *ENTR1* (Table 3, UR: *P*_*FDR*_ = 3.81 × 10^−27^, UE: *P*_*FDR*_ = 1.66 × 10^−24^). The ENTR1 protein is involved in cellular transport of cargo proteins from the endosome to the Golgi apparatus or for degradation in the lysosome (McGough et al., 2014) and has been suggested to play a role in cytokinesis (Hagemann et al., 2013). In a study where ENTR1 protein expression was blocked by RNA interference, there was a decrease in solute carrier family 2 member 1 glucose transporter protein (SLC2A1; previously known as GLUT1) (McGough et al., 2014). When examining whether this was due to increased SLC2A1 degradation, there was no evidence of increased transport of SLC2A1 to the lysosome. It was hypothesised that the decrease in SLC2A1 was mediated through regulation of transcription by ENTR1. SLC2A1 is responsible for approximately 30−40% of the glucose uptake in skeletal muscle, with the remainder transported through GLUT4 (Zisman et al., 2000; Rudich et al., 2003). As opposed to GLUT4 which is primarily expressed in skeletal muscle, SLC2A1 is widely expressed and is highly expressed on erythrocyte membranes (Krook et al., 2004). The control of SLC2A1 by ENTR1 in the context of the equine athlete is intriguing to speculate since SLC2A1 is expressed within equine lamellar tissue, and its expression is increased in hyperinsulinemia, therefore may play a role in the pathophysiology of equine laminitis (Campolo et al., 2016). *SLC2A1* is also differentially expressed in response to hypoxia, this has also been shown in equine chondrocytes *in vitro* after exposure to cobalt chloride (to mimic hypoxia) and in chondrocytes from osteoarthritis cases (Peansukmanee et al., 2009).

The most significant *trans* association in the UR cohort was between BIEC2-526896 and expression of the DEAH-box helicase 16 gene (*DHX16*) (Table 4). *DHX16* is an RNA helicase and is involved in regulation of translation and pre-mRNA splicing (Gencheva et al., 2010; Putiri and Pelegri, 2011). The gene located closest to BIEC2-526896 is the olfactory receptor family 12 subfamily D member 3 gene (*OR12D3*) with the TSS located 96.5 kb from the SNP. However, the zinc finger protein 311 gene (ZNF311) also relatively close to BIEC2-526896 (792.5 kb)(Consortium, 2017). *ZNF311* has previously been associated with telomere length in heterozygous ataxia-talengiectasia mutated (*ATM*) gene patients (Renault et al., 2017). As a member of the a krueppel c2h2-type zinc-finger protein family it is likely a transcription factor and has been associated with Biological Processes such as ‘regulation of transcription, DNA templated’ and ‘regulation of transcription by RNA polymerase II’(Consortium, 2017). The *trans* association between BIEC2-526896 and *DHX16* expression may therefore be mediated via the gene regulatory function of *ZNF311*.

The most significant *trans*-eQTL in the UE cohort was UKUL2765 and expression of the methylcrotonoyl-CoA carboxylase 2 gene (*MCCC2*) (Table 4). *MCCC2* encodes a subunit of 3-methylcrotonyl-CoA carboxylase (MCC), an enzyme which catabolises leucine (Stadler et al., 2005). Mutations within *MCCC2* have been found to result in MCC deficiency, which has varying implications for patients from no symptoms at all to death in early infancy (Fonseca et al., 2016). To date studies have yet to discern mutations which result in more or less severe disease phenotypes (Gallardo et al., 2001; Stadler et al., 2006). In terms of muscle physiology, *MCCC2* has been shown to be highly expressed in skeletal muscle of the red seabream fish (*Pagrus major*), which is likely due to high levels of protein metabolism within skeletal muscle (Abe et al., 2004). The TSS of the jumonji domain containing 4 gene (*JMJD4*) is located 71 bp from UKUL2765. The JMJD4 protein catalyses the hydroxylation of translation termination factor eRF1 lysine 63, which in turn enables the correct termination of translation and maintenance of translational fidelity (Feng et al., 2014). It is possible that the variation proximal to *JMJD4* is influencing expression of *JMJD4*, in turn altering expression of *MCCC2*. However, from the data available only one significant *cis*-eQTL for *JMJD4* was detected in the UR cohort and this was BIEC2-277622 located 257.8 kb downstream of the TSS (*P*_FDR_ = 6.58 × 10^−5^). Therefore it is not clear if UKUL2765 is tagging variation influencing *JMJD4* expression and mediating its influence on *MCCC2* through *JMJD4*.

Examination of eGenes previously shown to be involved in the skeletal muscle transcriptional response to exercise and training demonstrated that *TAL2* exhibited the most significant *trans*-eQTL in the UR cohort (BIEC2-658237; Table 6) and that this *trans*-eQTL was also highly significant in the UE cohort (*P*_FDR_ = 9.80 × 10^−8^; Table 4). *TAL2* encodes a basic-helix-loop-helix transcription factor (Xia et al., 1991; Langlands et al., 1997). Deletion of *TAL2* in mice has been shown to cause severe disruption of the development of the central nervous system, with new-born mice dying shortly post-partum (Bucher et al., 2000). *TAL2* has been shown to be vital for the development of gamma-aminobutyric acid (GABA, inhibitory neurotransmitter) signalling neurons in the developing midbrain, showing highly regulated and coordinated expression (Achim et al., 2013). When expression of *TAL2* was inhibited, neurons more closely resembled an excitatory glutamatergic phenotype (Achim et al., 2013). In terms of application in racing performance, GABA has previously been used as a calming agent in Thoroughbred racehorses, although it was banned from use in 2012. The GABA type A receptor associated protein like 1 gene (*GABARAPL1*) was also identified as a key regulator in the skeletal muscle transcriptional response to exercise (Bryan et al., 2017). In addition, we have previously reported functional enrichment of pathways related to neurodegenerative disorders in the transcriptional response to exercise (Bryan et al., 2017). Given the role of *TAL2* in GABAergic neuronal fate, this suggests a potential role for *TAL2* in the coordination of the response to exercise. These results suggest that the role of genes associated with neuronal differentiation and disease in the context of muscle and exercise warrants further investigation.

To identify common biological functions within genes identified under *cis* or *trans* regulation, enrichment analysis of Biological Processes among the gene sets was performed. Among the *cis* eGenes detected in both the UR and UE cohort, as well as *trans* eGenes in the UR cohort there was significant enrichment of cofactor metabolic processes (GO:0051186, Table S1, S3, S4). Genes within this cluster were related to metabolism and substrate utilisation, including vitamin and mineral binding and synthesis such as: selenium (selenium binding protein 1 gene, *SELENBP1*; and selenoprotein T gene, *SELENOT*), molybdenum (molybdenum cofactor sulfurase gene, *MOCOS*) and thiamine (thiamine triphosphatase gene, *THTPA*). Consequently, variation in the expression of genes associated with nutrient binding may lead to variation in the ability of horses to utilise such nutrients. In this regard, abundance of selenoprotein gene transcripts has been used to identify dietary requirements for selenium in rats and turkeys (Barnes et al., 2009; Taylor and Sunde, 2017). Given the inter-animal variation in expression observed for genes relevant to substrate binding, it may be possible to use this information to evaluate nutrient requirements for individual horses, or whether expression of these genes can be modulated through diet.

Many of these genes have also been shown to have functions relevant to exercise, and variation within the expression of these genes may underpin variation in athletic performance. For example, the selenium binding protein 1 gene (*SELENBP1*) is significantly downregulated in response to exercise (log_2_FC = −0.56; *P*_FDR_ = 3.71 × 10^−11^) (Bryan et al., 2017). In both normal and cancerous human cells *SELENBP1* has been shown to be highly variable in expression (Yang and Sytkowski, 1998). Functionally, the SELENBP1 protein has been shown to be involved in many cellular processes including detoxification (Ishii et al., 1996), cytoskeletal outgrowth (Miyaguchi, 2004) and regulation of reduction and oxidation within the cell (Jamba et al., 1997). *SELENBP1* was found be differentially expressed in blood in response to administration of human recombinant erythropoietin in human endurance athletes (Durussel et al., 2016; Wang et al., 2017), suggesting a potential role in haematopoiesis and its regulation. In the UE cohort; the DDB1 and CUL4 associated factor 12 (*DCAF12*) and guanosine monophosphate reductase (*GMPR*) genes both exhibited significant *cis*-eQTL (*DCAF12*: *P*_FDR_ = 0.02; *GMPR*: *P*_FDR_ = 4.17 × 10^−3^). These genes, in addition to *SELENBP1*, were also shown by Wang et al. (2017) to be differentially expressed in blood in response to human recombinant erythropoietin. Variation in the expression of these genes may therefore potentially underpin variation in haematological phenotypes in horses, which may in turn influence traits relevant to aerobic capacity. It is also noteworthy that selenium deficiency has been associated with significant myopathy (White muscle disease) (Lofstedt, 1997; Delesalle et al., 2017) and reduced exercise tolerance in horses (Brady et al., 1978; Avellini et al., 1999). In addition, selenoproteins have been shown to be involved in several metabolic pathways and the response to oxidative stress in muscle (Rederstorff et al., 2006). These findings suggest an important role for selenium, and its associated biochemical machinery, in the correct functioning of skeletal muscle and muscle metabolism. This highlights the importance of selenium in the context of exercise and provides a potential role for variation in expression of genes relevant to selenium metabolism in determining metabolic function within the muscle.

The cofactor metabolic process cluster also contained genes relevant to mitochondrial function and oxidative phosphorylation. These included genes within the coenzyme Q synthesis pathway: coenzyme Q3 hydroxylase (*COQ3*), coenzyme Q7 (*COQ7*) and coenzyme Q8A (*COQ8A*). The coenzyme Q complex is a critical component of the electron transport chain during oxidative phosphorylation, moving electrons from complexes I and II to complex III (Lenaz and De Santis, 1985; Turunen et al., 2004; Stefely and Pagliarini, 2017). COQ7 and COQ8A are required for coenzyme Q biosynthesis (Mollet et al., 2008; Stefely et al., 2016). Human patients with *COQ8A* mutations suffered seizures and other neurological symptoms and showed reduced coenzyme Q within skeletal muscle (Jacobsen et al.; Mollet et al., 2008). An eQTL for *COQ8A* in skeletal muscle has already been identified in horses, with a 227 bp SINE insertion in the promotor region of *MSTN* (g.66495326_66495327ins227) on ECA18 associated with increased expression of *COQ8A* (previously known as *ADCK3*) in Thoroughbreds (Rooney et al., 2017). However it should be noted that this increase in COQ8A expression did not appear to accompany an COQ8A protein abundance, with no difference in COQ8A protein abundance across genotypes (Rooney et al., 2017). This may be due to COQ8A having a regulatory role in coenzyme Q biosynthesis (Acosta et al., 2016). Electron transport chain complex activity assays, as well as assays using the exogenous application of ubiquinone, suggested a difference in the abundance of coenzyme Q across genotypes at this locus (Rooney et al., 2017). Suggesting variation at this SINE insertion is associated with *COQ8A* expression as well as coenzyme Q abundance. Therefore eQTL in the current study associated with *COQ8A*, and indeed other genes within the coenzyme Q biosynthetic pathway, may result in variation in synthesis of the coenzyme Q complex and have downstream implications for mitochondrial function.

We have for the first time systematically catalogued eQTL in equine skeletal muscle, both at rest and post-exercise. Previous investigations of eQTL in skeletal muscle have focussed primarily on human T2D (Sharma et al., 2011; Keildson et al., 2014a; Sajuthi et al., 2016a; Langefeld et al., 2018)and meat quality traits in production animals (Ponsuksili et al., 2015; Gonzalez-Prendes et al., 2017; Pampouille et al., 2018; Gonzalez-Prendes et al., 2019; Velez-Irizarry et al., 2019). Our investigation of eQTL in the context of skeletal muscle and exercise present some of the only work to-date in this area (Kelly et al., 2014). These results provide novel information concerning the regulation of gene expression in horses and can provide a framework for interpreting future GWA studies of athletic and performance traits in Thoroughbreds. While conventional differential expression analyses can identify genes up- and downregulated in response to an exercise stimulus, they cannot detect inter-individual variation in the transcriptional response. The response to exercise and training has been shown to be highly heritable in humans (Timmons et al., 2010; Bouchard et al., 2011) and it is also likely to be highly heritable in horses. Therefore, systems genetics approaches that integrate differential gene expression with genome variation represent an excellent strategy for dissecting the genetic architecture of these complex anatomical and physiological traits.

## Data accessibility statement

Data are available on request from the UCD Technology Transfer Office for researchers who meet the criteria for access to confidential data.

## Author contributions

GF performed computations and functional analyses. KB and PAM assisted in analysis and pipeline development. CLM, KFG and GF performed biopsy sample collections. JAB and GF prepared RNA. GF wrote the manuscript in close consultation with EWH, LMK and DEM. All authors were involved in study design, implementation of the research and preparation of the manuscript.

## Funding

This research was funded by Science Foundation Ireland (SFI/11/PI/1166 and 18/TIDA/6019).

## Conflicts of interest statement

EWH, DEM and LMK are named inventors on multiple international patents relating to the application of *MSTN* variation in the prediction of race distance performance; none of which is relevant to the data/results reported in this manuscript.

## Acknowledgments

We would like to thank J.S. Bolger for access to his horses, and staff at Glebe House stables for their assistance, particularly B. O’Connor and P. O’Donovan. This research was conducted with the financial support of Science Foundation Ireland (grant no. SFI/11/PI/1166 and 18/TIDA/6019).

